# Hydrogel array patterning using 3D-printed microfluidic inserts to control cell-cell and cell-ECM interactions

**DOI:** 10.64898/2025.12.21.695682

**Authors:** Matthew D. Poskus, Mohammadmahdi Eskandarisani, Ioannis K. Zervantonakis

## Abstract

By shaping biochemical gradients and extracellular matrix cues within the local microenvironment, cellular spatial organization plays a critical role in regulating tissue development, homeostasis, and disease progression. Microfluidic platforms are highly suitable for the study of these cell-cell and cell-matrix interactions as they precisely control cell arrangement and gradients compared to conventional experimental systems. Cells are often embedded within hydrogels to improve physiological relevance by enabling matrix-mediated signaling. However, many designs restrict the number and arrangement of hydrogels or generate gradients in only one dimension, limiting their ability to recapitulate complex tissue architectures. To address this need, we introduce a 3D printed microfluidic insert compatible with microplates that allows patterning of up to ten unique hydrogel arrays in two dimensions and generation of parallel or orthogonal concentration gradients. We first develop a physics-based computational model of hydrogel filling to define design parameters that ensure robust hydrogel patterning. We then establish perpendicular concentration gradients on timescales relevant to biological experiments. Furthermore, we demonstrate high cell viability in our 3D-printed devices and control of fibroblast migration across multiple patterned hydrogels. Finally, we monitor the recruitment of primary human monocyte towards cell-free and fibroblast-seeded 3D collagen matrices. Our microfluidic insert platform is compatible with high-throughput automation workflows and allows for interrogation of spatially variant signals that regulate cell migration and cell-cell signaling in physiologically-relevant 3D microenvironments.

## Introduction

Cell behavior is tightly coupled to spatial context. Through direct contact and exchange of secreted factors, cells continuously integrate biochemical and biophysical cues from their neighboring cells and the extracellular matrix (ECM) to coordinate development and maintain tissue homeostasis^1,2^. Disruptions to this organization of cell-cell and cell-ECM interactions are often hallmarks of disease^2,3^. These interactions occur over tens of microns for juxtacrine signaling and hundreds of microns for paracrine signaling to shape local microenvironments^4^. Cell positioning and gradients are also interdependent; cells secrete and consume factors that establish concentration gradients to regulate critical cellular functions, including cell migration, which in turn influence tissue structure^5,6^. Defining the role of spatial organization and gradients in regulating cell behavior is essential for elucidating processes that underlie both normal tissue function and chronic diseases^3,7^.

*In vitro* systems that recapitulate tissue architecture can interrogate these interactions under controlled conditions. Conventional plate-based culture platforms lack the three-dimensional (3D) organization and soluble-factor gradients characteristic of native tissues, limiting their ability to capture and model spatially regulated signaling. Microfluidic platforms address these constraints by leveraging micron-scale features to position cells with high precision and produce concentration gradients, enabling the quantitative study of cell-cell interactions at the single-cell level^8,9^. When combined with 3D hydrogels that mimic key aspects of ECM, these organ-on-a-chip systems better approximate tissue environments^10,11^ and have become powerful platforms for modeling organ-level interactions in physiologically relevant conditions^12–16^. A widely used strategy for hydrogel compartmentalization is capillary pinning, wherein surface tension forces at engineered microstructures halt meniscus advancement during hydrogel loading to create barrier-free interfaces after hydrogel gelation^17^. This phenomenon depends on the feature geometry and material surface wettability to prevent spilling of the hydrogel under the pressure of loading via pipette. The applied pressure increases the apparent contact angle until a critical advancing contact angle threshold is achieved and the interface advances^18^.

Most microfluidic platforms with a 3D ECM are configured as a single hydrogel compartment bordered by culture medium channels or additional hydrogel channels^19–23^. These channels may be supplemented with soluble factors or cells to generate biochemical gradients useful for studies of directed cell migration^23,24^. With this design, patterning and gradients are limited to a single dominant linear or radial dimension, thereby limiting the ability to recreate complex tissue architectures or study response to multidirectional stimuli. Additional designs permit patterning of arbitrary hydrogel shapes using surface tension^25–27^; however, these designs introduce space between patterned gels that disrupt ECM signaling as cells transition between regions, a critical process in multicellular organization or migration studies. New approaches for patterning hydrogels are needed to overcome these limitations.

3D printing overcomes barriers associated with conventional photolithography techniques for microfluidic fabrication. Vat photopolymerization 3D printing enables layer-by-layer curing of a photosensitive resin to produce molds for polydimethylsiloxane (PDMS) devices or directly print devices for cell culture^28,29^. The latter approach is low cost, supports rapid prototyping, and enables fabrication of geometries that are difficult to achieve with PDMS-based replica molding^30,31^. However, resin biocompatibility must be considered as cytotoxic resin compounds can leach into cell culture medium^29,32^. The biocompatible polymer parylene-c can be applied via chemical vapor deposition to improve cytocompatibility of 3D-printed devices^33–35^. This layer acts as a barrier to resin compounds due to its low chemical permability^36^.

Here, we present 3D-printed, microplate-compatible microfluidic inserts that enable multidimensional hydrogel patterning and gradient formation within a continuous extracellular matrix. Our platform supports patterning of up to ten distinct hydrogels (nine primary and one secondary) and four integrated medium reservoirs allow generation of orthogonal or opposing biomolecular gradients. We first investigated the impact of two critical design parameters: hydrogel gap height and plate substrate wettability on capillary pinning using a computational model and validated these findings experimentally. Next, we demonstrated the ability to sustain orthogonal concentration gradients over several days. After validating the biocompatibility of our device, we assessed the 3D migration of human fibroblasts subjected to biochemical stimulation and infiltration of primary human monocytes into 3D hydrogels seeded with fibroblasts. Our microfluidic inserts will enable future investigation of cell-cell signaling mechanisms not accessible with one-dimensional patterning systems and can be directly integrated with high-throughput well-plate systems.

## Materials and Methods

### Microfluidic insert fabrication

Microfluidic devices were designed in AutoDesk Inventor Professional 2024 (Autodesk, USA) and prepared for printing using Chitubox (Chitubox, China) slicing software. A Phrozen Sonic Mini 8K liquid crystal display printer (Phrozen Technology, Taiwan) was used to fabricate inserts in Liqcreate Bio-Med Clear resin (Liqcreate, Netherlands) with an exposure time of five seconds and 25μm layer height. Following printing, devices were submerged in fresh isopropyl alcohol (Fisher Scientific, USA) and underwent three cycles of cleaning, where one cycle consists of washing in an Anycubic Wash and Cure 2.0 station (Anycubic, China) for 30 minutes then processing for five minutes in an ultrasonic cleaner (Kaimashi, China). Inserts were postcured in an Anycubic Wash and Cure 2.0 station for 3 hours. Parylene-c coating was performed using a LABCOTER 2 (Specialty Coating Systems, USA) using 10g DPX-C precursor. System temperature parameters were preconfigured to a furnace temperature of 690°C, chamber temperature of 135°C, and vaporizer temperature of 175°C. Prior to culture with cells, devices were sterilized in 70% ethanol for 30 minutes, rinsed with water twice, then allowed to air dry in a sterile environment. Devices were inserted into one of four 24-well microplates: Glass bottom (cat.nr P24-1.5H-N, Cellvis, USA), untreated polystyrene (cat.nr 3738, Corning, USA) and tissue culture treated (cat.nr, 662165 Greiner, Germany) plates with or without plasma treatment to increase wettability. To create plasma-treated plates, tissue culture treated microplates were exposed to plasma generated at 2 Torr using a plasma cleaner (Harrick Plasma, USA) for 120s immediately prior to use.

### Collagen hydrogel preparation and device filling

Primary and secondary hydrogels were prepared at a concentration of 2mg/mL using buffered rat tail collagen type I (Corning, USA) and stored on ice. An electronic pipette (BRAND, USA) was used to dispense 2µL of collagen solution into the nine primary hydrogel regions at the lowest dispense speed. Unused microplate wells were filled with water to minimize evaporation prior to incubation at 37°C and 5% CO2 for 25 minutes to polymerize the hydrogel. The secondary hydrogels were filled until the meniscus extended to the edges of the hydrogel region using approximately 1µL of solution. Secondary hydrogel ports were filled in a clockwise pattern to minimize risk of bubble trapping. Devices were incubated for 20 minutes and subsequently filled with 75µL cell culture medium in each of the four medium reservoirs.

### Fluorescent bead assay

Collagen hydrogels containing yellow-green or red fluorescent 1µm microspheres (2% aqueous suspension, Invitrogen, USA) were added at a concentration of 1.5µL microspheres per 200µL hydrogel solution. Hydrogels were filled following the protocol outlined above into glass bottom, untreated, tissue culture treated and plasma-treated plates. First, the green bead hydrogel was injected into the primary hydrogel ports, and the red bead hydrogel was injected using the secondary hydrogel ports. A wide-field Nikon Ti2 microscope (Nikon, Japan) was used to capture fluorescence images of the beads following polymerization. Z-stacks were acquired using an ImageXpress Micro Confocal (Molecular Devices, USA) at a height interval of 10µm for a total height of 400µm. 3D renderings were generated using the ImageJ 3D Viewer plugin.

### Contact angle measurements

Surfaces were cleaned using 70% ethanol and wiped with a lint-free wipe to remove debris prior to and between each measurement. Microplates were modified to access a cross section of the well with a camera. A mixture of deionized water plus a few drops of food dye was used to enhance visibility. A 2µL droplet of dyed water was carefully pipetted onto the surface of the plate and images were acquired using a portable camera mounted to a stand. Quantification was performed in ImageJ by measuring the angle between the substrate and droplet at the three-phase contact line.

### Capillary pinning numerical model

3D simulations were performed in COMSOL Multiphysics 6.2 (COMSOL Multiphysics, Sweden) using Two-Phase Flow, Phase Field module to model the dynamics of the air-liquid interface. The liquid hydrogel precursor phase was modeled using water. The simulation domain consisted of a single primary hydrogel region and symmetry boundary conditions were applied to reduce the computational domain to an eighth of the full model. This decision was based on the insert design that has four identical outlets and symmetric geometry. A constant volumetric inflow rate was set to 3.5µL/s determined by measuring the time required to dispense a fixed volume using the electronic pipette in experiments. A mesh and parameter refinement study was performed to determine the mesh discretization and phase field interface thickness. We focused on the transition between hydrogel pinning and mixing for contact angles between 40-50° and a gap height of 50μm. Simulation results converged as mesh size and interface thickness were reduced across the coarse, medium and fine settings.

### Dextran diffusion assay

Solutions of 125µg/mL10kDa Cy5 dextran (Thermo Fisher Scientific, USA) and of 100µg/mL 70kDa Texas red fluorescent dextran (Thermo Fisher Scientific, USA) were diluted in Dulbecco’s Modified Eagle Medium (DMEM) cell culture medium. 75µL of Cy5 dextran solution or Texas red solution was added to the topmost or rightmost reservoir of the insert, respectively. The two remaining reservoirs were filled with 75µL cell culture medium. Unused wells were filled with phosphate buffered saline (PBS) to minimize evaporation, and the plate was sealed with parafilm. Timelapse images were acquired using a Cytation C10 (Agilent, USA) microscope and devices were incubated at 37°C for the duration of the experiment. The fluorescent intensity was calculated within a small area (approximately 400µm diameter circle) in the center of each hydrogel region.

### Cell culture

Fibroblasts were passaged at 70% confluency and maintained at 37°C and 5% CO2 using DMEM containing 10% (v/v) heat-inactivated fetal bovine serum (FBS) and 1% (v/v) penicillin/streptomycin. Cells were trypsinized using 0.05% Trypsin. Human peripheral blood mononuclear cells (PBMCs) were isolated from healthy donors, and pan monocytes were isolated using a pan monocyte isolation kit #130-096-537 (Miltenyi Biotec, Germany). Pan-monocytes were frozen after isolation and thawed before each experiment.

### Viability assay

Fibroblasts were incubated with a solution of 200nM CellTracker Green CMFDA diluted in DMEM cell culture medium for 30 minutes and washed twice with PBS prior to trypsinization. A buffered solution of 2mg/mL collagen type I hydrogel (Corning, USA) was prepared containing fibroblasts at a concentration of 1.5M cells/mL and dispensed into the devices. After hydrogel polymerization, 75µL of DMEM cell culture medium (10% FBS) containing 50nM SYTOX Deep Red was added to each of the four device reservoirs, to monitor dead cells. Images were acquired using an ImageXpress Micro Confocal microscope (Molecular Devices, USA) in the GFP and Cy5 channel every day for four days. Cells were maintained at 37°C and 5% CO2. Viability was calculated using Cellprofiler^37^ as the ratio of live cells (GFP+ minus Cy5+ area) in each field divided by the total cells (GFP+ area).

### Fibroblast migration assay

Fibroblasts were incubated with a solution of 200nM CellTracker Green CMFDA diluted in DMEM cell culture medium for 30 minutes and washed twice with PBS prior to trypsinization. The primary hydrogel solution was prepared by mixing a buffered solution of 2mg/mL collagen type I hydrogel (Corning, USA) containing fibroblasts at a concentration of 1.5M cells/mL and dispensed into the devices. After hydrogel polymerization, 75µL of DMEM cell culture medium containing either 0% FBS (control) or 10% FBS was added to all medium reservoirs. Images were acquired using an ImageXpress Micro Confocal microscope (Molecular Devices, USA) in the GFP and Cy5 channel every day for three days. The number of migrating fibroblasts was calculated using Cellprofiler^37^ and a custom MATLAB (Mathworks, USA) script. The number of migrating cells was calculated as the number of cells within the secondary hydrogel region at each timepoint by drawing regions of interest to exclude cells in the primary regions.

### Monocyte infiltration assay

Monocytes were incubated with a solution of 200nM CellTracker Green CMFDA diluted in RPMI cell culture medium, respectively, for 30 minutes and washed with PBS prior to trypsinization. Two buffered solutions of 2mg/mL collagen type I hydrogel (Corning, USA) were prepared containing either no fibroblasts or fibroblasts at a concentration of 1.5M cells/mL to be dispensed into the devices as the primary hydrogel. Following polymerization of the hydrogel, pan-monocytes were mixed with a solution of 50% RPMI, 50% DMEM (v/v) at a density of 0.5M cells/mL and 75µL of this solution was added to each reservoir of devices. Images were acquired with an ImageXpress Micro Confocal (Molecular Devices, USA) in the GFP channel every day for three days. Quantification was performed using Cellprofiler^37^ and a custom MATLAB script. Regions of interest were drawn around each of the nine primary hydrogel regions. The total number of pan-monocytes in all regions were summed for each timepoint.

### Statistical Analysis

Statistical analysis was performed using Prism Graphpad. Unless otherwise specified, statistics were conducted using paired multiple t test for biological replicates.

## Results

### Continuous hydrogel array patterning in 3D-printed, well-plate embedded microfluidic inserts

We developed a 3D-printed cell culture platform that is placed directly into standard 24-well microplates for high-throughput studies (**Figure 1A-B**). The inserts feature external grooves that plastically deform when pressed in the well to prevent movement during hydrogel or cell culture medium loading. This design eliminates laborious device preparation steps common to protocols for PDMS-based microfluidics, such as punching filling ports and binding to a cover-glass. In addition, it enables greater design freedom compared to photolithography, which requires alignment and assembly of multiple photoresist layers for complex geometries.

**Figure 1:**
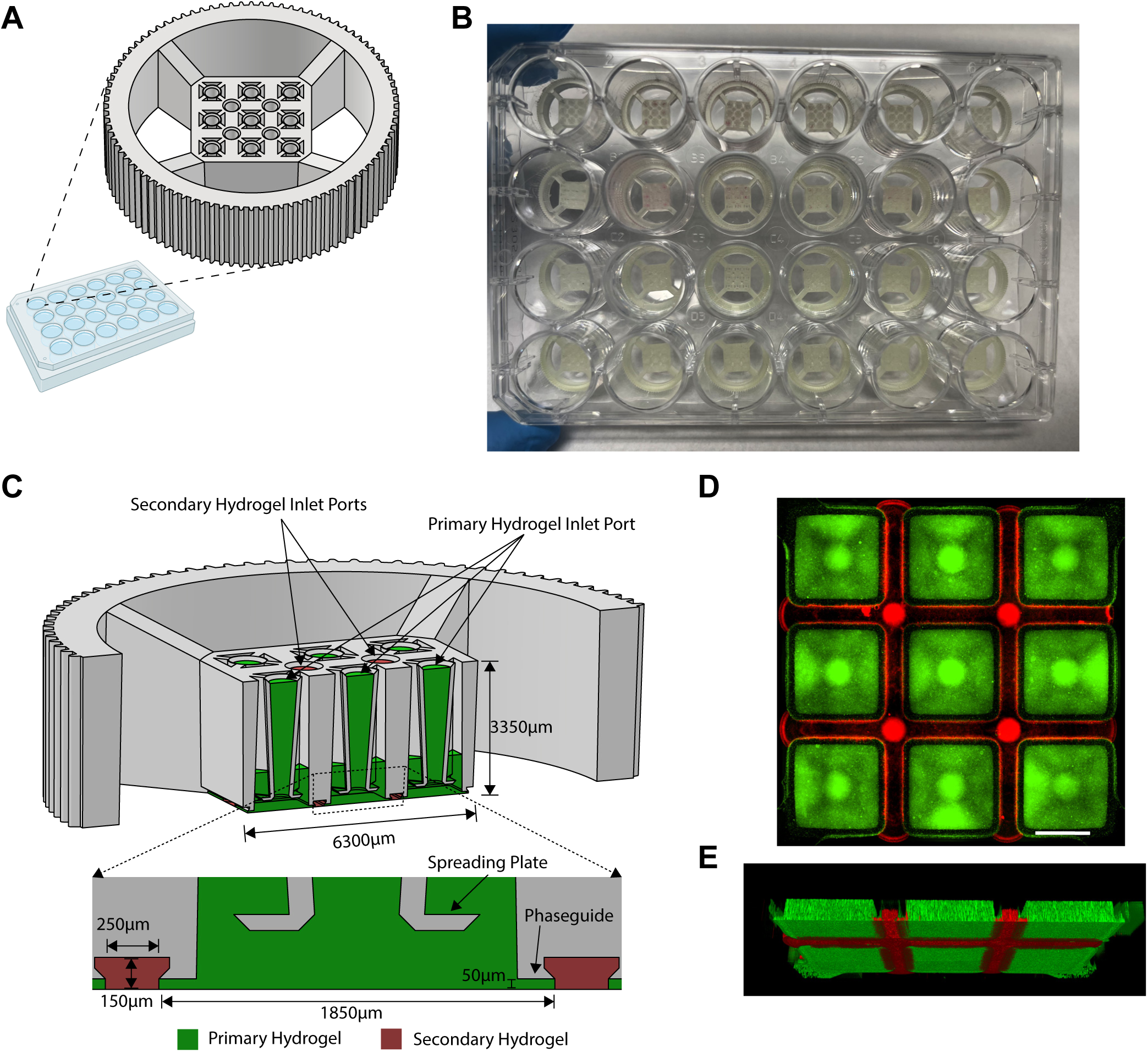
Microfluidic insert design for two-dimensional hydrogel patterning. (A) Well-plate compatible design concept. (B) Inserts fitted into a standard 24-well microplate. (C) CAD model cross section view of microfluidic insert and hydrogels. Bottom insert shows the spreading plate and phase guide features that ensure confinement of the primary (green) hydrogel while it is filled from the nine inlet ports. The secondary hydrogel (red) is filled from the remaining four inlet ports and occupies the empty space between adjacent primary hydrogels. (D) Experimental demonstration of patterned primary (green) and secondary (red) hydrogels mixed with fluorescent beads using 3D-printed inserts placed on a tissue culture treated plate. Image was acquired with an inverted widefield fluorescent microscope at the focal plane of the well-plate bottom. Scalebar: 1000µm. (E) 3D confocal rendering to visualize the volume of patterned hydrogels (a volume of 400µm was imaged).

The hydrogel patterning area consists of nine independent primary hydrogel regions arranged in a three-by-three grid separated by a narrow secondary hydrogel region (**Figure 1C**). The hydrogels are filled via pipette through inlet ports located at the top of the insert. Briefly, the primary hydrogels are first filled and polymerized in an incubator, followed by addition and polymerization of the secondary hydrogels. A phaseguide at the border of the primary hydrogel regions confines the primary hydrogels to prevent hydrogel mixing, and any excess hydrogel during filling flows upward through the concentric outlets. The result is a continuous hydrogel domain comprised of up to ten unique primary and secondary hydrogels (**Figure 1D-E**).

### Computational modeling and experimental validation to investigate how gap height and plate substrate hydrophilicity determine hydrogel patterning success

The capillary pinning effect occurs at the primary hydrogel phaseguide and is dependent on the hydrophilicity of the surfaces in contact with the hydrogel precursor air-liquid interface. The liquid at this interface contacts the phaseguide and bottom plate substrate with contact angles θ*_p_* and θ_s_, respectively. Pressure from hydrogel filling increases these contact angles according to the Young-Laplace Equation^38^, which relates the pressure sustained by the interface to geometrical parameters and contact angles. The height of the hydrogel in this region is bounded by bottom plate surface and phaseguide and this gap height is small relative to the width of the pillar. The resulting curvature of the meniscus is dominated by this height and can be approximated as flow through two parallel plates, for which the relationship between the pressure sustained by the interface, materials hydrophilicity, and geometry is given by^39^:

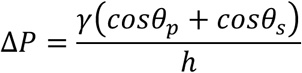

where *ΔP* is the pressure drop across the air-liquid interface, γ is the surface tension coefficient, and *h* is the gap height. The phaseguide is coated with parylene-c and therefore the contact angle θ*_p_* is fixed; however, the gap height and substrate contact angle θ_s_ can be varied by changing the device geometry or plate substrate material. Therefore, we sought to investigate how these two parameters affect hydrogel pinning to ensure robust hydrogel patterning for all plate types (**Figure 2A**). A wide range of plate materials with various surface treatments are available for specific applications. For example, high content screening assays may require high optical clarity glass-bottom plates or functionalization to improve cell adhesion for culture. We measured water contact angles on parylene-c coated devices and on the following microplate surfaces: untreated polystyrene, tissue culture (TC) treated polystyrene, and glass bottom plates, with a plasma-treated TC microplate used as positive control (**Figure 2B**).

**Figure 2:**
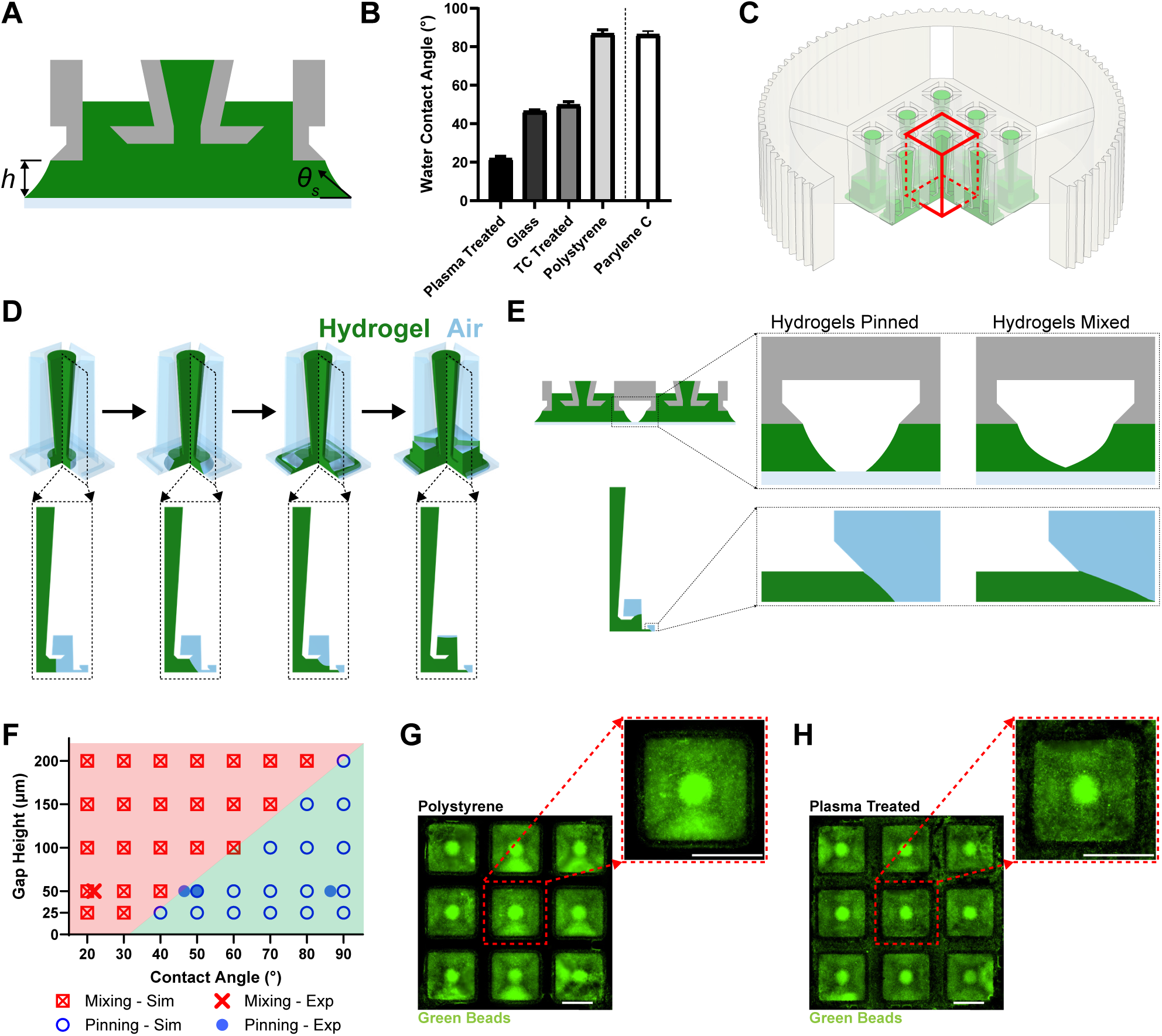
Hydrogel patterning is dependent on device geometry and plate substrate wettability. (A) Schematic of critical design parameters: gap height and substrate contact angle. (B) Water contact angle measurements for four plate substrates and parylene-c coated devices. Plate substrates include glass bottom, untreated polystyrene, tissue culture (TC) treated polystyrene, and plasma-treated TC. (C) Computational domain consisting of a single primary hydrogel region is three-dimensional and outlined with a box. (D) Simulated dynamics of hydrogel filling for a gap height of 50µm and substrate contact angle of 50°. As the hydrogel (green) fills the region, air (light blue) is displaced. Top row shows the 3D simulation domain outlined in panel D and bottom row shows cross-section. (E) Hydrogel pinning criterion for simulations physically represents contact between two adjacent hydrogels that results in interfacial mixing. (F) Comparison of simulated (open symbols) and experimental (filled symbols) hydrogel pinning results for a range of gap heights and substrate contact angles. Hydrogel mixing is outlined with a cross and hydrogel pinning with a circle. (G-H) Successful hydrogel pinning for untreated polystyrene plate (gap height 50μm and contact angle: 90°) or hydrogel mixing in plasma-treated TC plates (gap height 50μm and contact angle: 20°). Red rectangles show a single primary hydrogel region as shown in the simulations (panel D). Scalebar: 1000µm.

We sought to identify which gap height would be sufficient to constrain hydrogels for all plate types. Toward this end, we developed a computational model of filling dynamics of a single hydrogel domain for a range of gap heights (25-200µm) and substrate contact angles based on our wettability measurements (20-90°) (**Figure 2C-D**). We considered the hydrogel successfully confined if the hydrogel did not exceed the boundaries of the computational domain (primary hydrogel region) for the duration of the simulation. Physically, this extension would result in contact between two adjacent primary hydrogels during filling, resulting in both mixing and occupation of the secondary hydrogel channel (**Figure 2E**). Our simulation results revealed that more hydrophobic substrates (untreated polystyrene) sustain pinning at greater gap heights compared to hydrophilic surfaces (TC-treated polystyrene and glass bottom) with lesser contact angles (**Figure 2F**). As the 50µm gap was the maximum height predicted to successfully confine hydrogels for all three standard microplate materials, we fabricated devices with this height and validated these findings experimentally (**Figure 2F-G**). Furthermore, our model predicted failure to confine hydrogels on the plasma-treated plate. Hence, we filled devices inserted into a plasma-treated plate and confirmed that the primary hydrogels were not confined, spilling into the secondary hydrogel region and mixing (**Figure 2H**). Given that the 50µm height was able to confine hydrogels for all plates, this height was selected for all future experiments.

### Generation of orthogonal gradients across patterned hydrogels

We next evaluated the capability to generate multidirectional concentration gradients of two sizes of fluorescent dextran in our device. We first filled the primary and secondary hydrogel space with a 2mg/mL collagen type I hydrogel. PBS containing either 10kDa Cy5 or 70kDa Texas Red fluorescent dextran was added to the top or right medium reservoirs, respectively. PBS was added to the remaining reservoirs to establish orthogonal left-right and top-bottom gradients of these molecules (**Figure 3A**). We monitored these devices using confocal microscopy for 4 days (**Figures 3B-C**) and quantified the fluorescence intensity in three primary hydrogel regions (close, center, far) in the middle row (70kDa dextran) or middle column (10kDa dextran) (**Figure 3D-E**). The fluorescence intensity increased over time for all regions. As expected, a faster increase in intensity was observed in the close region compared to the central and far region.

**Figure 3:**
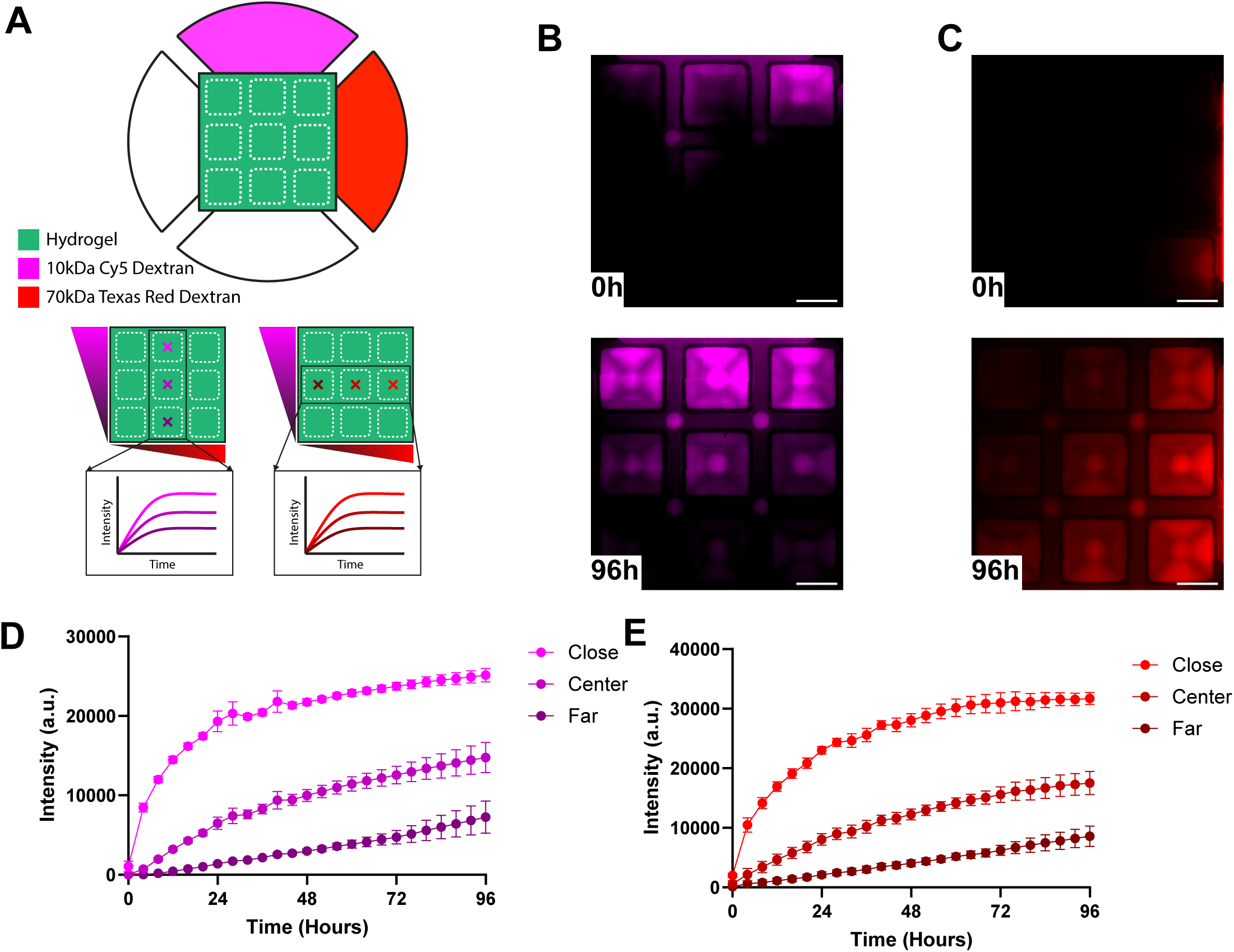
Two-directional gradient generation within hydrogel arrays. (A) Orthogonal concentration gradients are formed by filling adjacent reservoirs with 10kDa (magenta, top reservoir) or 70kDa (red, right reservoir) fluorescent dextran solution in 3D-printed inserts placed on a tissue culture treated plate. (B) Representative images of 10kDa Cy5 dextran or (C) 70kDa Texas red dextran gradients at indicated timepoints. Scalebar: 1000µm. (D) Quantification of the 10kDa or (E) 70kDa dextran intensity profiles in the center of three primary hydrogel regions shown with crossed in panel (A). Error bars are representative of standard error of the mean of at least four devices.

### 3D-printed microfluidic inserts preserve cell viability and enable monitoring of fibroblast migration and monocyte infiltration in three-dimensional collagen matrices

We next sought to highlight the utility of patterning arrays of hydrogels that form a continuous ECM network by monitoring the migration of cells between hydrogel regions and the recruitment of cells from the device reservoirs into the hydrogels. First, we confirmed that cell viability is sustained in our parylene coated microfluidic inserts by monitoring death of human fibroblasts cultured within a 3D collagen type I matrix over four days. Fibroblasts were seeded into the primary hydrogels and exhibited a spindle morphology after initial seeding (**Figure 4A**). High cell viability (>90%) in the parylene-coated devices was maintained for four days and was comparable to viability found in control collagen droplets seeded directly in microplates without 3D-printed inserts (**Figure 4B**).

**Figure 4:**
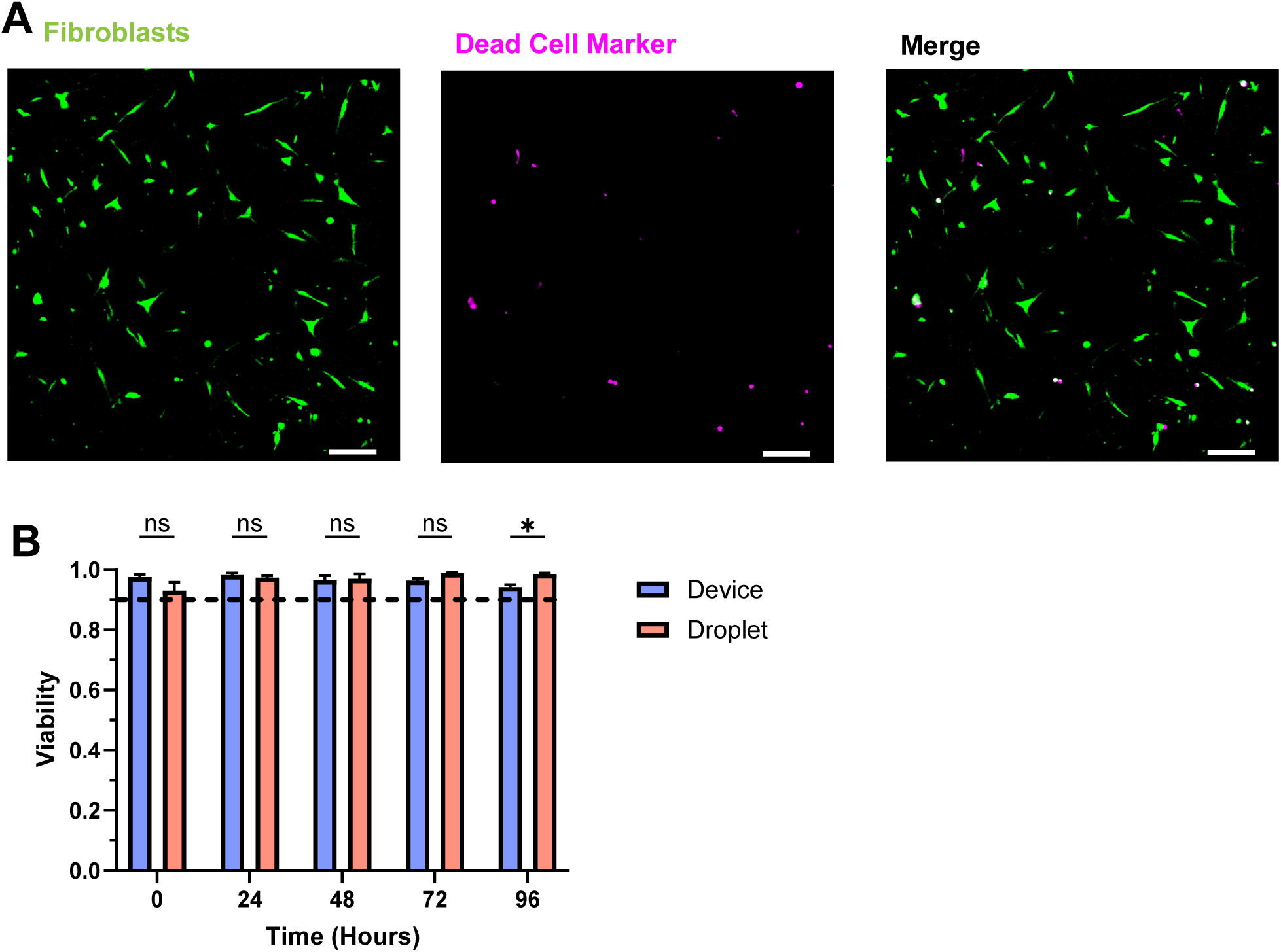
Parylene coated 3D-printed microfluidic inserts maintain cell viability. (A) Representative images of CellTracker Green CMFDA stained fibroblasts incubated with SYTOX Deep Red after four days of 3D culture in 3D-printed inserts placed on a tissue culture treated plate. Scalebar: 100µm. (B) Quantification of fibroblast viability within 3D-printed inserts on tissue culture treated plates (blue bar) or within macroscale collagen droplets on tissue culture treated plated (red bars, no 3D-printed insert). Statistical significance was assessed using paired t tests between groups for each timepoint. * indicates p<0.05.

Having established the biocompatibility of our 3D-printed inserts, we next measured the migration of fibroblasts from the primary hydrogel to the secondary hydrogel in the presence or absence of serum. We seeded fibroblasts into all nine primary collagen type I hydrogels surrounded by a cell-free secondary collagen type I hydrogel and measured the number of cells in each region for three days under conditions of starvation medium (0% FBS) or complete medium (10% FBS) (**Figure 5A**). In both conditions, fibroblasts migrated into the secondary gels (**Figures 5B-C**); however, the magnitude of the migration differed between conditions. As expected, in the absence of serum fewer fibroblasts migrated into the secondary hydrogel compared to complete medium after 72 hours (**Figure 5D**).

**Figure 5:**
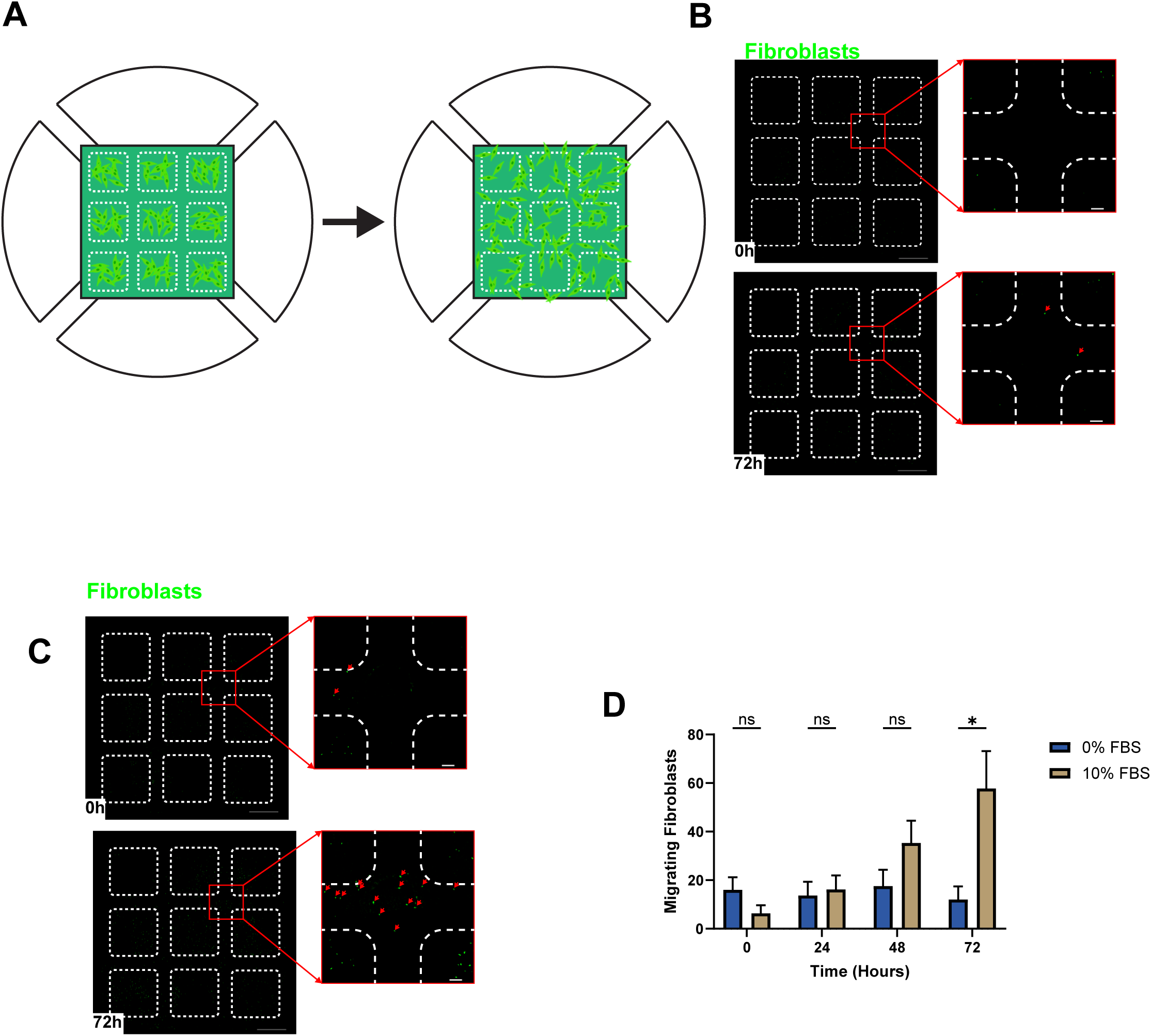
3D fibroblast migration between hydrogel regions under biochemical stimulation. (A) Schematic of experimental setup to monitor fibroblast migration from primary to secondary hydrogels. Both primary and secondary hydrogels are formed using 2mg/ml collagen type I. (B-C) Representative images of fibroblast migration in 0% or 10% FBS conditions. 3D-printed inserts placed on a tissue culture treated plate. Red arrows show fibroblasts. (D) Quantification of the number of fibroblasts that migrated into the secondary hydrogel. Scalebar: 1000µm or 100µm for inset images. Statistical significance was assessed using paired t tests between groups for each timepoint. * indicates p<0.05.

We also evaluated the capacity of fibroblasts to recruit monocytes into collagen type I matrices. Our experimental design included (a) fibroblast seeding into the primary hydrogel space separated by a cell-free secondary hydrogel and (b) cell-free primary/secondary hydrogels (**Figure 6A**). Monocytes were suspended in cell culture medium and added to the cell culture medium reservoirs. The number of monocytes infiltrating into the primary hydrogels was measured every day for two days under conditions of monocyte coculture with fibroblasts and monocyte monoculture (**Figure 6B&C**). Compared to cell-free primary hydrogels, the addition of fibroblasts to the primary hydrogels increased the number of monocytes recruited to this region after 48 hours (**Figure 6D**).

**Figure 6:**
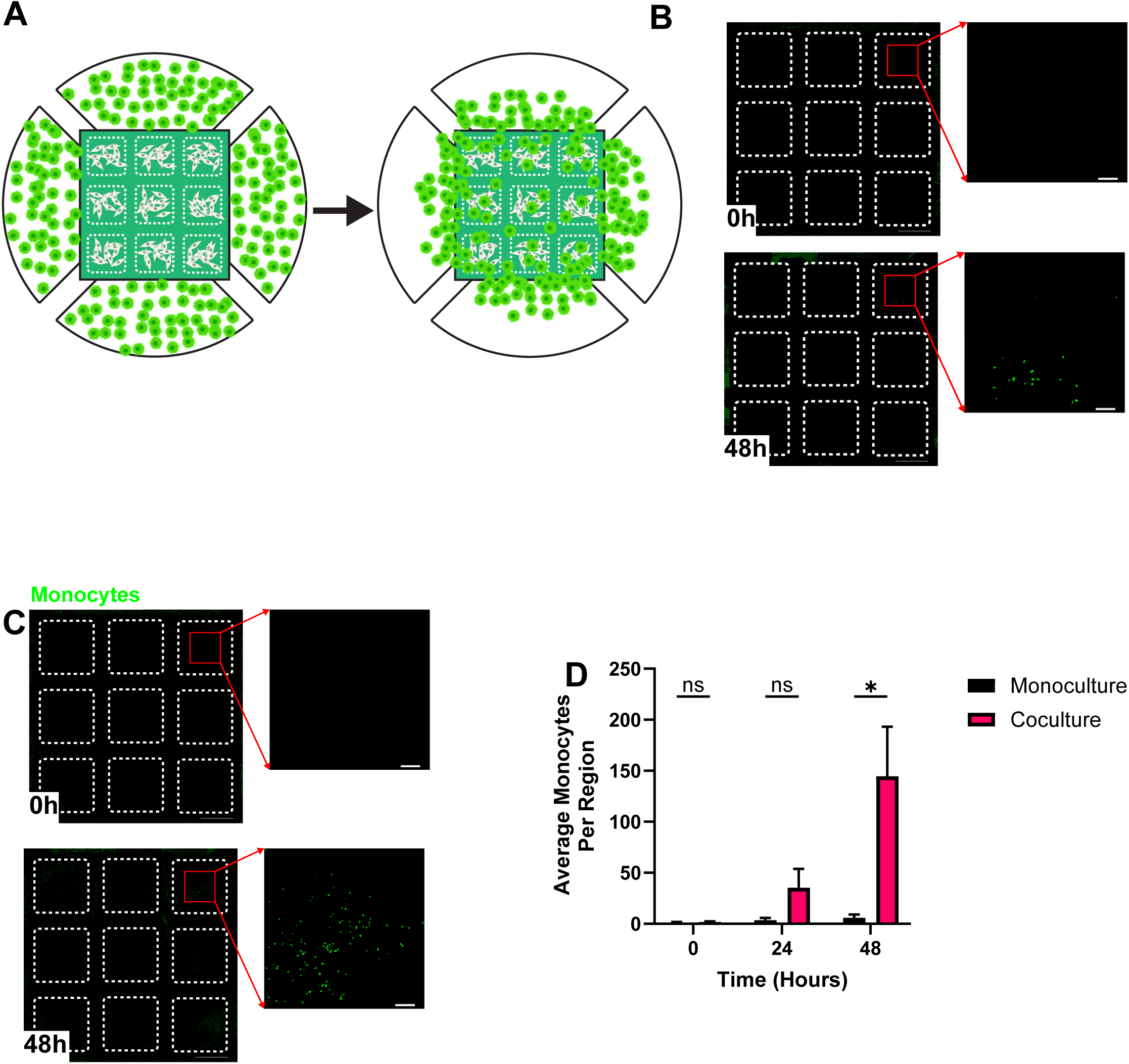
Fibroblast-driven primary human monocyte infiltration into 3D collagen hydrogels. (A) Schematic of experimental setup to monitor fibroblast-driven 3D recruitment of monocytes. Both primary and secondary hydrogels are formed using 2mg/ml collagen type I. (B) Representative images of monocyte recruitment within collagen type I primary hydrogels after 48 hours in monoculture or (C) coculture with fibroblasts. 3D-printed inserts placed on a tissue culture treated plate. (D) Quantification of the total number of monocytes in primary hydrogels over time. Scalebar: 1000µm or 100µm for inset images. Statistical significance was assessed using paired t tests between groups for each timepoint. * indicates p<0.05.

## Discussion

Signaling between cells is inextricably linked to spatial proximity, as cells communicate through both direct contact and secreted factors. Existing microfluidic platforms fall short of reproducing the complex tissue architectures of extracellular matrix heterogeneity present *in vivo*. To address this gap, we developed a 3D-printed microfluidic insert that integrates with microplates and enables spatial control of cell-cell and cell-ECM interactions in a 3D matrix. The device design features an array of 3D hydrogels and medium reservoirs that enables two-dimensional control of hydrogel placement and generation of orthogonal biochemical gradients. By arranging the patterned hydrogels adjacent to one another, a continuous matrix is formed that preserves cell-ECM signaling as cells transition between regions. We used a computational model to explore how gap height and substrate hydrophilicity influence capillary pinning across common microplate materials and then validated these predictions experimentally. After confirming high cell viability, we applied the platform to measure fibroblast migration and assess monocyte recruitment by fibroblasts within 3D collagen matrices.

Our findings align with prior work examining surface wettability, fibroblast motility, and fibroblast-immune cell interactions. We first measured water contact angles of relevant substrates as inputs to our capillary pinning computational model. The wettability of parylene-c measured here was consistent with previous reports^34,40^ and trends across substrate contact angles agree with similar studies showing glass plates exhibited greater hydrophilicity than untreated polystyrene^21^. Simulations revealed that capillary pinning depends on gap height and substrate wettability, in agreement with earlier studies^21,25^. We next showed that fibroblast viability is maintained in the parylene-c coated devices, which is consistent with previous reports^33–35^. As a functional demonstration of our system, we quantified fibroblast migration in the presence or absence of fetal bovine serum and found that serum promotes the motility of fibroblasts in 3D, consistent with prior work^41^. In the same platform, fibroblasts recruited primary human monocytes into the 3D matrix via soluble factors. Fibroblasts are known to produce cytokines and growth factors that attract monocytes in diverse contexts^42,43^.

Capillary pinning is a precise, convenient hydrogel patterning technique owing to its dependence on readily tunable geometrical and material parameters. Prior studies have used this approach to generate hydrogel arrays that vary in number and size of regions to probe cell-cell interactions^19–22,26,27^. These hydrogels can be placed adjacent to each other^19–22^, or separated by cell culture medium channels^26,27^. In the latter configuration, cells migrating between regions must exit the matrix, traverse a 2D substrate, then re-enter a hydrogel. This discontinuity at the channel-hydrogel boundary disrupts matrix-mediated cellular signaling in cell-cell and cell-ECM interactions studies. To date, the largest reported hydrogel arrays consist of five adjacent hydrogels^21^ or seven hydrogels separated by cell culture medium^26^. In contrast, our platform supports up to ten adjacent hydrogels that collectively form a continuous matrix, thereby expanding patterning capabilities while maintaining more physiological conditions. The dimensions of each region determine the spatial resolution of cell patterning, with smaller regions enabling finer control over cell positioning. Most devices emphasize a 1D, single dominant patterning dimension, where characteristic sizes range between 0.6-2mm^19–21^. Our insert instead offers a 2D array with hydrogel regions of comparable scale (2.1 mm), enabling flexible patterning within a standard microplate format.

The patterning capabilities of this platform could be extended by increasing the quantity of patterning regions and spatial resolution by capitalizing on advances in laboratory automation and 3D printing technologies. The grid-like configuration of our hydrogel array is scalable to greater numbers of hydrogel regions, but practical use is restricted by the time required for manual pipetting relative to the gelation time, particularly when working with multiple devices at once. Laboratory automation equipment could alleviate these constraints, as exemplified in a commercially available microfluidic platform that can be paired with an open source pipetting robot^44^. In parallel, ongoing improvements in 3D printing technologies offer finer feature resolution, where efforts have focused on reducing printer pixel size and optimizing resin formulation. Smaller pixel sizes improve XY resolution, thus reducing the minimum printable feature size^45^. In this work, we used a consumer-grade 3D printer with 22μm pixels, whereas recent custom systems and specialized commercial printer are capable of pixel sizes of 2-7.6μm^46,47^. Similarly, resin chemistry can be tailored to the printer optics to increase print resolution^48,49^. For example, smaller internal channels can be printed by increasing the concentration of photo absorber in the resin, which reduces the penetration depth of light^49^. When custom resins and printers are combined and optimized, features on the order of tens of microns can be fabricated^49,50^, compared with the hundreds of microns size typical of many commercial printers and resins^29,51^. These efforts highlight improvements in print fidelity, and by extension, enable finer hydrogel patterning control that can further enhance the spatial resolution of our platform.

## Acknowledgements

The authors acknowledge grant support from the US National Institutes of Health (R35GM150815 to I.K.Z.).

